# Nitrogen enrichment alters selection on rhizobial genes

**DOI:** 10.1101/2024.11.25.625319

**Authors:** Caleb A. Hill, John G. McMullen, Jay T. Lennon

## Abstract

1

Mutualisms evolve over time when individuals belonging to different species derive fitness benefits through the exchange of resources and services. Although prevalent in natural and managed ecosystems, mutualisms can be destabilized by environmental fluctuations that alter the costs and benefits of maintaining the symbiosis. In the rhizobia-legume mutualism, bacteria provide reduced nitrogen to the host plant in exchange for photosynthates that support bacterial metabolism. However, this relationship can be disrupted by the addition of external nitrogen sources to the soil, such as fertilizers. While the molecular mechanisms underpinning the rhizobia-legume symbiosis are well-characterized, the genome-wide fitness effects of nitrogen enrichment on symbiotic rhizobia are less clear. Here, we inoculated a randomly barcoded transposon-site sequencing (RB-TnSeq) library of the bacterium *Ensifer* (*Sinorhizobium*) *meliloti* into soils containing a host plant, alfalfa (*Medicago sativa*), under conditions of low and high nitrogen availability. Although plant performance remained robust to fertilization, nitrogen enrichment altered gene fitness for specific traits and functions in the rhizobial partner. Genes involved in carbohydrate metabolism showed increased fitness irrespective of soil nutrient content, whereas fitness gains in quorum-sensing genes were only observed in high-nitrogen environments. We also documented reductions in the fitness of nucleotide metabolism and cell-growth genes, while genes from oxidative phosphorylation and various amino-acid biosynthesis pathways were detrimental to fitness under elevated soil nitrogen, underscoring the complex trade-offs in rhizobial responses to nutrient enrichment. Our experimental functional genomics approach identified gene functions and pathways across all *E. meliloti* replicons that may be associated with the disruption of an agronomically important mutualism.

**Importance:** Understanding the evolutionary dynamics of the rhizobia-legume mutualism is important for elucidating how plant-soil-microbe interactions operate in natural and managed ecosystems. Legumes constitute a significant portion of global food production and generate 25% of all terrestrially fixed nitrogen. The application of chemical fertilizers can disrupt the mutualism by altering the selective pressures experienced by symbiotic rhizobia, potentially affecting gene fitness throughout the microbial genome and leading to the evolution of less productive or cooperative mutualists. To investigate how exogenous nitrogen inputs influence gene fitness during the complex rhizobial lifecycle, we used a barcoded genome-wide mutagenesis screen to quantify gene-level fitness across the rhizobial genome during symbiosis and identify metabolic functions affected by nitrogen enrichment. Our findings provide genomic insight into potential eco-evolutionary mechanisms by which symbioses are maintained or degraded over time in response to changing environmental conditions.

## 3 Introduction

Soil microorganisms can form beneficial symbiotic relationships with plants that allow them to withstand environmental fluctuations across space and time. These mutualistic interactions typically involve the reciprocal exchange of resources or metabolites that would otherwise be inaccessible to each partner [1]. However, there are costs to maintaining mutualisms, and these associations can be disrupted by changing environmental conditions. In the rhizobia-legume symbiosis, plants secrete chemoattractive flavonoids that aid in the recruitment of rhizobia from the soil habitat. Infection through root hairs leads to the intracellular uptake of rhizobia and development of endosymbiotic root nodules [2]. While some rhizobia live normally within nodules, others differentiate into a specialized bacteroid cell-type which fixes atmospheric nitrogen N_2_ within the low-O_2_ nodule environment, supplying hosts with a limiting nutrient (NH_4_) in exchange for photosynthates. This relationship is costly and complex, with different metabolic requirements and selective challenges at each stage [3]. For example, *Rhizobium leguminosarum* requires a set of *>* 600 essential genes to nodulate and fix nitrogen [4]. Meanwhile, legumes supply organic carbon to rhizobia nodules which could otherwise be used for plant maintenance and growth [5]. The symbiosis can be destabilized by nitrogen enrichment [1, 6], which leads to host divestment from mutualism in favor of inorganic soil nitrogen uptake. Evolutionarily, fertilization can relax selection for genes that are involved in symbiosis and downregulate the maintenance of existing nodules [1, 6, 7].

While the molecular mechanisms underlying the rhizobia-legume symbiosis are well-characterized, it is less clear how nitrogen fertilization alters mutualistic processes from a functional genomics perspective. In many rhizobial strains, essential nitrogen-fixing machinery is located on horizontally transferred symbiotic plasmids [2]. Prolonged nitrogen enrichment selects for mutations on these plasmids, resulting in less efficient and less cooperative symbiotic relationships [6]. To better understand how nitrogen enrichment affects whole-genome selection on rhizobia genes and functions across symbiotic stages, we inoculated alfalfa (*Medicago sativa*) plants with a randomly barcoded transposon-site sequencing library (RB-TnSeq) of their symbiont, *Ensifer* (*Sinorhizobium meliloti*), in soils maintained under low vs. high nitrogen conditions. RB-TnSeq employs barcoded transposons to generate whole-genome knockout libraries, enabling the quantification of individual mutant lineages through barcode sequencing [8, 9]. This allows for the assessment of genome-wide gene fitness by comparing changes in barcode relative abundances before and after growth under specific experimental conditions [8, 9]. We uncovered genes and metabolic pathways under selection at low and high nitrogen in undifferentiated and bacteroid cell-types. Our functional genomics approached allowed for the characterization of a complex microbial symbiosis and the identification of mechanisms by which mutualism can be destabilized by nitrogen fertilization.

### 3.1 Nitrogen enrichment altered rhizobia-legume interactions

Although plant growth was not affected by nitrogen enrichment (ANOVA: *F*_(1,8)_ = 0.27, *P* = 0.62, *η*^2^ = 0.033, [Fig S1]), we documented compositional differences in the recovery of barcoded *Ensifer* mutants that were associated with alfalfa grown in low vs. high nitrogen (PERMANOVA, *R*^2^ = 0.12, *F*_(1,10)_ = 1.69, *P* = 0.003). These genome-wide differences also existed between individual pots (PERMANOVA, *R*^2^ = 0.11, *F*_(1,10)_ = 1.55, *P* = 0.033). While genome-wide differences between undifferentiated and bacteroid cell-types were not significant (PERMANOVA, *R*^2^ = 0.091, *F*_(2,10)_ = 0.66, *P* = 0.98), we did observe a significant shift in the distribution of fitness effects (DFEs) in mutants extracted from bacteroids at low vs high nitrogen (two-sample Kolmogorov-Smirnov with false-discovery rate [FDR] adjustment, *D* = 0.04, *q* = 0.0004; [Fig. S2]). Together, these patterns suggest that rhizobia in nodules experienced altered host and environmental selection for particular functions, a hypothesis we explored further using functional enrichment analysis (Figs. 1, 2).

**Fig 1.**
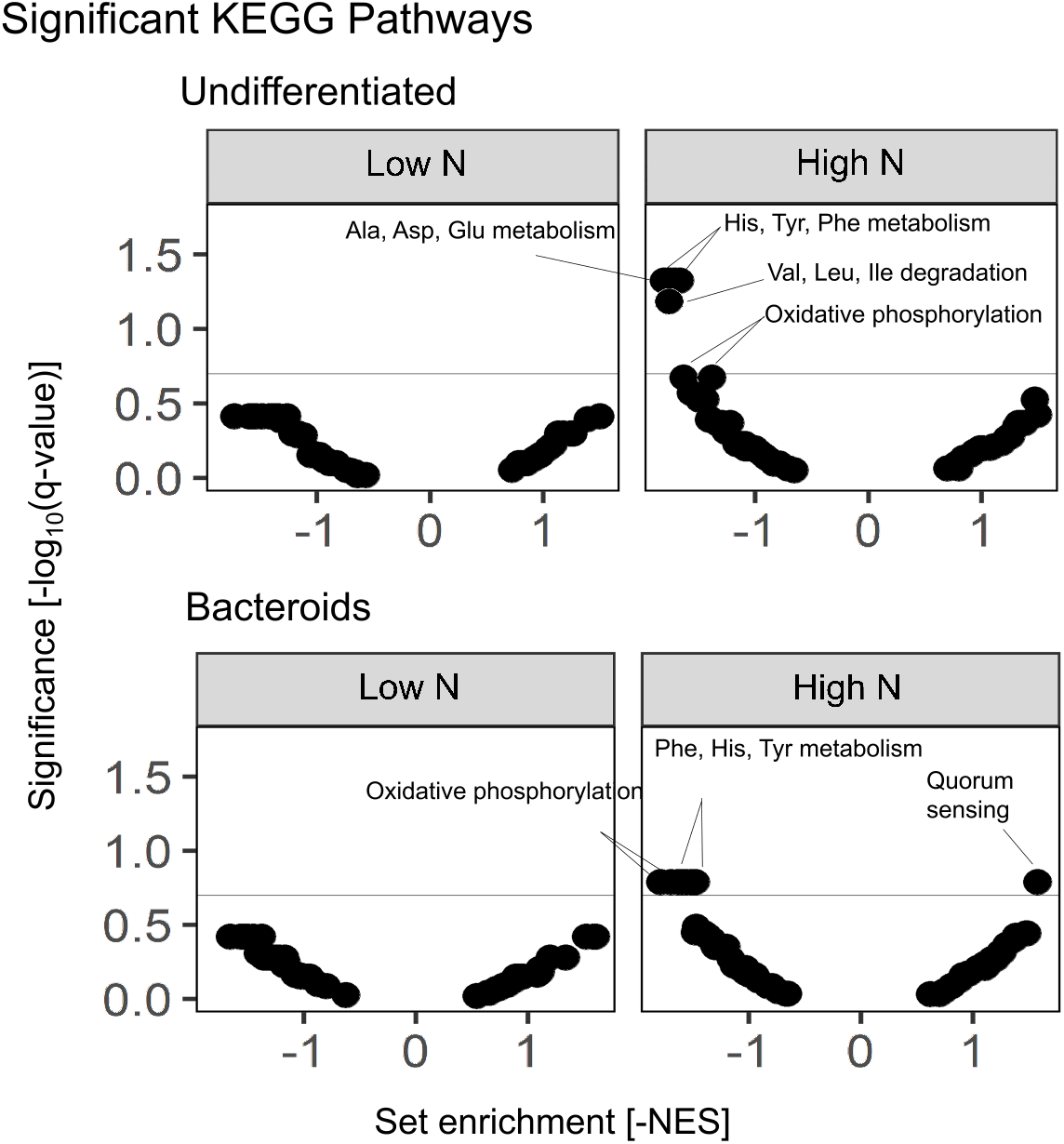
Gene-set enrichment of KEGG categories by nitrogen level and cell-type. Nitrogen enrichment altered selection on metabolic functions in both undifferentiated nodule bacteria and bacteroids. We ranked genes by fitness using a modified *t* -statistic calculated from the raw fitness scores and queried KEGG pathway annotations against them using gene-set enrichment analysis (GSEA). The x-axis depicts set enrichment score, which is obtained by multiplying the “normalized enrichment score” from GSEA by -1 [-NES]. If the set enrichment score is positive, then genes from that set are beneficial to fitness. That is, RB-TnSeq mutants with knockouts for genes in that set grew less effectively during the experimental period and were depleted in the endpoint pool. If the set enrichment score is negative, then genes from that set are detrimental to fitness. That is, mutants with knockouts in that set grew more effectively and were enriched in relative abundance at the end of the experiment. The y-axis depicts statistical significance for that gene set’s score after false discovery rate adjustment (FDR), with the FDR threshold of 0.2 depicted by the horizontal line. Analyses were separately conducted on genes annotated as essential vs. nonessential for symbiosis; only one set was significant within the nonessential genes for either cell-type. This was “Val, Ile, Leu degradation” in the high-nitrogen nodule bacteria, a pathway which some studies have found to be important for symbiosis [10, 11] and may have been mis-annotated by our reciprocal best-hits method.

**Fig 2.**
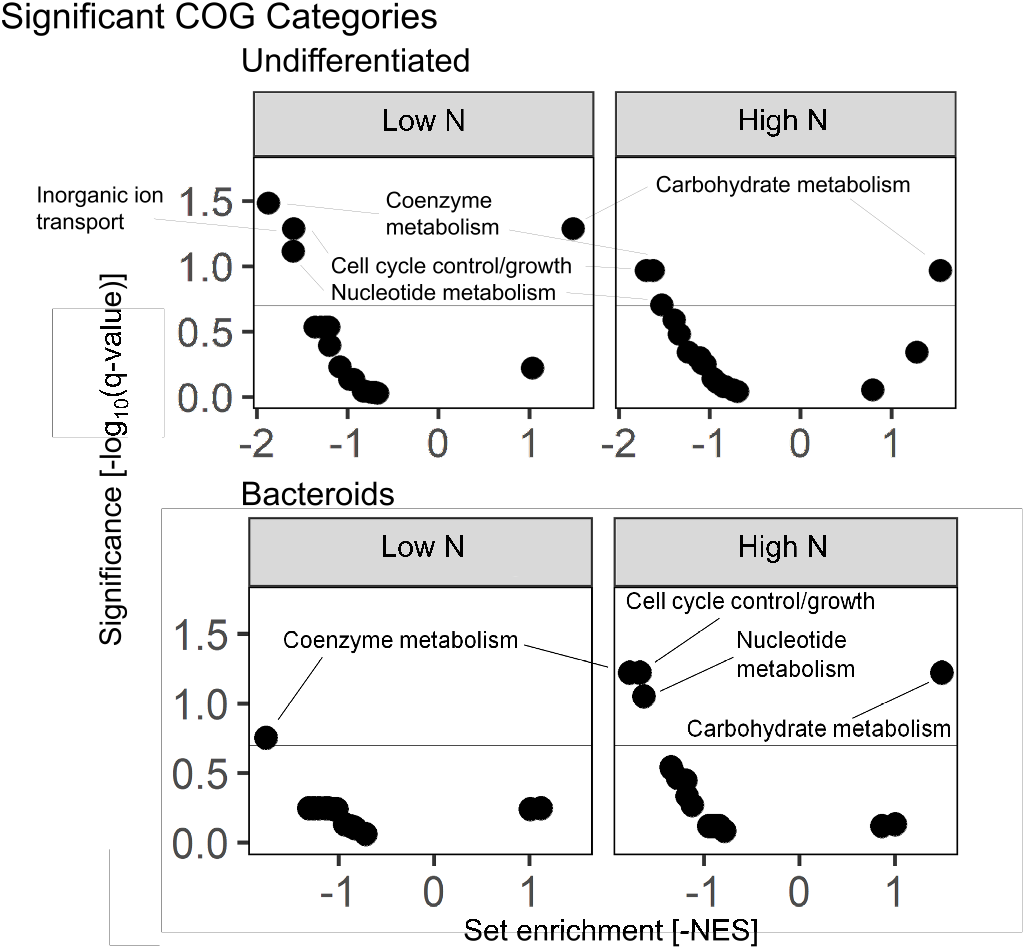
Gene-set enrichment analysis of COG categories by nitrogen level and cell-type. Some metabolic functions experienced selection at both low and high nitrogen, though this effect was stronger among undifferentiated nodule bacteria. We ranked genes by fitness using a modified *t* -statistic calculated from the raw fitness scores and queried COG categories against them using gene-set enrichment analysis (GSEA). The x-axis depicts set enrichment score, which is obtained by multiplying the “normalized enrichment score” from GSEA by -1 [-NES]. As described in Fig. 1, if the set enrichment score is positive, then genes from that set are beneficial to fitness. If the set enrichment score is negative, then genes from that set are detrimental to fitness. The y-axis depicts statistical significance for that gene set’s score after false discovery rate adjustment (FDR), with the FDR threshold of 0.2 marked with a horizontal line. Analyses were separately conducted on genes annotated as essential or nonessential for symbiosis, but there were no significant sets among non-symbiosis genes, so all results presented here represent sets of symbiosis genes.

### 3.2 Relaxed selection under elevated nitrogen

Using a gene-set enrichment analysis (GSEA) defined by KEGG pathways, we identified genes and functions that experienced altered selection under contrasting soil nitrogen across undifferentiated and bacteroid subpopulations (Fig. 1). For example, nitrogen enrichment had a negative effect on the fitness of certain amino-acid biosynthesis genes in both groups (Fig. 1). Previous studies have shown that high biosynthesis of histidine, tyrosine, and branched-chain amino-acids improves nodule competitiveness in other rhizobia-legume systems [4, 10, 11]. Negative fitness of these biosynthetic functions in our experiment are consistent with reduced competition for establishment and nodulation under high nitrogen. This effect cascades from undifferentiated cells to bacteroids, as the latter are derived from the former (Fig. 1). Similarly, cytochrome *c* biosynthesis genes exhibited low fitness under high nitrogen conditions, likely reflecting the advantage rhizobia gained from the reduced burden of maintaining costly metabolic machinery. Additionally, quorum-sensing genes showed positive fitness exclusively in bacteroids under high nitrogen conditions (Fig. 1). This finding suggests that QS-regulated processes may be important for bacteroid formation and/or survival in high-nitrogen environments [12, 13]. Together, our findings demonstrate that increased nitrogen may relax selection for nodule competitiveness by favoring reduced growth, given that competitive establishment and nodulation depend on amino-acid biosynthetic capacity.

### 3.3 General environmental effects on gene fitness

Beyond nitrogen-dependent fitness effects, we identified changes in gene fitness across nitrogen levels that may be related to general influences of the rhizosphere environment and demands of within-nodule metabolism. Using a GSEA based on coarse-grained metabolic functions (i.e., COG categories), we found evidence for increased fitness across nitrogen treatments and cell-types for carbohydrate metabolism genes, including gluconeogenic enzymes, beta-glucan synthesis genes, and myo-inositol catabolism genes. Our findings are consistent with previous reports that these functions are important for persistence in the alfalfa rhizosphere across free-living and rhizospheric niches [2, 10]. Additionally, myo-inositol catabolism is required for rhizopine signaling, an important nutritional mediator between undifferentiated nodule bacteria and the host plant [14]. Genes involved in cell growth such as peptidoglycan biosynthesis (*cpoB*) and chromosomal partitioning (*scpA*/*scpB*) (Table S2) returned negative fitness scores across treatments and cell-types, as did nucleotide and amino-acid metabolism genes such as the rate-limiting enzyme for *de novo* guanine synthesis (*guaB*), purine and ribonucleotide genes (*pyrB* /*pyrC* /*pyrE* /*pyrF, purC* /*purF* /*purK* /*purN* /*purQ*), and threonine biosynthesis gene (*thrB*) (Table S2). These growth-related genes are expressed at a high level by undifferentiated nodule bacteria as they undergo multiple rounds of endoreduplication during their terminal differentiation into bacteroids [15]. Mutants which fail to endoreduplicate will not terminally differentiate or fix nitrogen, instead continuing to grow and divide within the nodule. These mutants exploit the benefits of symbiosis without incurring the associated energetic or reproductive costs, effectively functioning as uncooperative mutualists. Nitrogen-dependent effects are likely present among bacteroids in this analysis due to low mutant recovery within one of the high-nitrogen bacteroid replicates, indicating high attrition of bacteroids at high nitrogen (Table S1). This pattern may also arise due to founder effects and arrested growth within the smaller bacteroid subpopulation. These results starkly contrast with our pathway-level analysis which found no metabolic pathways under selection at low nitrogen (Fig. 1), indicating that nitrogen enrichment disrupts specific symbiotic processes on top of general pressures of the rhizosphere and nodule environments.

## 4 Conclusion

Our experiment demonstrates how RB-TnSeq can be used to quantitatively understand how nitrogen enrichment alters selection on genes of a microbial symbiont. Our experiment revealed genome-wide patterns of selection for rhizobial gene function in association with alfalfa (*Medicago sativa*), complementing previous studies which have primarily focused on individual symbiotic genes and functions often encoded on symbiotic plasmids [6, 7]. In fact, we found that genes under selection at high nitrogen were significantly concentrated on the chromosome (Chi-square with Yates’ continuity correction, *χ*^2^ = 6.3, *df* = 1, *P* = 0.012). Our results suggest that nitrogen enrichment may relax selection for competitiveness by favoring mutations that slow or arrest biosynthesis (Figs. 1, 2). This corroborates findings from genome-scale metabolic modeling regarding the high biosynthetic demands of growth in the rhizosphere and the metabolic pressures of within-nodule growth such as myo-inositol catabolism and endoreduplication required for bacteroid differentiation [10, 14, 15]. Additionally, fitness differences between undifferentiated and bacteroid subpopulations suggest traits that favorably influence development to the bacteroid stage (Fig. 1). Most genes under selection relate to the underlying metabolic machinery that power the expression of core symbiosis genes (Table S2), demonstrating how disruption of mutualism can lead to whole-genome selection on fundamental physiological processes. Future RB-TnSeq experiments could expand our understanding of the eco-evolutionary dynamics of this symbiosis by studying how other independent variables — such as host genotype, phages, or interspecies/interkingdom interactions — affect microbial fitness at different nitrogen levels. By testing RB-TnSeq library growth across an array of environmental variables and habitats, we can better understand and predict the eco-evolutionary processes that influence whether microbial symbioses persist or dissipate, as well as how they function through time.

## 5 Acknowledgments

We acknowledge David Merritt and Doug Rusch at the Indiana University Center for Genomics and Bioinformatics for assistance with sequencing and data analysis; Morgan Price, Kelly Wetmore, and Adam Deutschbauer for providing us with the the Sm1021 mutant library used in this experiment; Valentine Trotter for advice on RB-TnSeq experimental protocol; Emmi Mueller for assistance in the laboratory; Jen Lau and Irene Newton for feedback on an earlier version of the manuscript. JGM contributed to this manuscript outside of their capacity at Bayer and the views and opinions of the authors do not necessarily reflect those of Bayer Crop Sciences.

## 7 Data Availability

All code used in this study is available at https://github.com/LennonLab/rhizo.rb.tnseq. RB-TnSeq data analysis scripts can be found on the FEBA codebase, and mutant pool information and results from previous RB-TnSeq experiments [8, 9] can be found here.

