## Supplementary Information for "Nitrogen enrichment alters selection on rhizobial genes"

### Methods

#### 1.1 Plant and microbial culturing

*Medicago sativa* seeds were obtained from Ernst Conservation Seeds and surface-sterilized by repeated washing: once in 6% sodium hypochlorite (NaClO) for 10 min, once in 70% ethanol (EtOH) for 10 min, and then three times with deionized water. Soil substrate was created by mixing Turface MVP calcined clay (Turface Athletics, LLC) with Metro-Mix 820 (Sun-Gro Horticulture, Ltd.) at a 2:1 ratio and autoclave-sterilizing at 121 °C for two 24 h cycles with at least 24 h rest in between. Pots were assembled in two tiers. We used a handheld power drill to make holes in the bottom-center of GA-7 Magenta boxes (Magenta, LLC), which were placed on top of undrilled boxes that servved as a water reservoir. Cotton wicks were cut (8 cm), burned at the ends to eliminate fraying, and run through the holes to create a wick-watering system that could be refilled with sterilized water by removing the lower unit. Approximately 300 mL of soil was put into the upper tier, taking care to run the wick through the middle of the soil. To wash soil of soluble nutrients and ensure equal distribution, the upper pot was removed from the reservoir and deionized water was run through soil multiple times. Pots were then sealed with manufacturer-provided lids and shaken vigorously to homogenize the soil. The reassembled two-tier pots were autoclaved for 90 min to resterilize the completed unit. The reservoir water was replaced prior to seed germination and whenever necessary using autoclaved deionized water.

After allowing sterile soil to cool, three small wells were made in equal spacing around the top of the wick, and 3-4 *M. sativa* seeds were aseptically transferred into each soil well and reburied. Seeds were germinated for one week, uncovered, and then culled to one plant per well to give three plants per pot. Plants were grown in a Percival PGC-40L2 (Percival Scientific, LLC) growth chamber with a 16:8 light:dark photoperiod at 50% relative humidity, 23 °C daytime and 18 °C nighttime temperature and a light intensity of 1200 $\mu$mol/m$^2$/s. Water reservoirs were covered with aluminum foil to prevent algal growth. Deionized water was autoclaved in large carboys for 90 min and let cool before use.

We cultivated *Ensifer* (*Sinorhizobium*) *meliloti* Sm1021 in batch culture in low-sodium LB (5g NaCl/L). We estimated cell density based on measurements of absorbance (OD600) converted to colony-forming units (CFU) with an OD-to-CFU linear regression. Bacteria were centrifuged for 3 min at 7,500 *g* to pellet cells before being resuspended in 500 µL phosphate buffered saline (PBS), pH = 7.4. We then added $1 \times 10^7$ CFU directly to the bases of *M. sativa* shoots along with PBS-inoculated negative controls.

After one week of growth to allow the infection and nodulation process to occur, half the plots were fertilized with aqueous ammonium nitrate. Nitrate fertilizer was added once per week for three weeks to ensure that high nitrogen levels were achieved without inducing fatal nitrogen toxicity in plants. To mix fertilizer, 0.97 g $NH_4NO_3$ was added to 40 mL of sterile e-pure water. We then pipetted 4 mL of liquid fertilizer evenly across the whole surface of each pot. This resulted in an $NH_4NO_3$ concentration of $11\,\mathrm{mg\,N/m^2}$ .

#### 1.2 Experimental setup

To assess the effect of nitrogen enrichment on gene-level fitness of symbiotic *S. meliloti*, 15 pots were created: 10 library-inoculated plants and five PBS-inoculated controls. Of the 10 library plants, half were fertilized with nitrogen, giving five replicates per treatment group. Plants were allowed to grow for eight weeks prior to harvesting at which point nodule samples were taken. PBS plants served as negative controls for the effect of symbiont presence/absence on plant growth. Once harvested, nodule samples were separated into samples enriched for undifferentiated populations inhabiting the nodule or for fully differentiated bacteroids living within the symbiosome, yielding another set of treatment groups for the purposes of sequencing and microbial genomics.

#### 1.3 Data collection

To harvest nodules, plants were cut from the base of the shoot and plant height was measured. To measure biomass, plants were cut to fit into pre-weighed paper bags, dried out for 72 h at 80 °C, and weighed again. To account for water loss from the bags, three empty bags were weighed, dried as described, endpoint masses were averaged to determine water loss per bag, and this value was subtracted from the final mass measurement. Plant data were pooled and averaged on a per-pot basis. Nodule count and biomass data were collected as well; aposymbiotic control plants were checked for nodules and a small number of failed nodules were putatively identified on three of the five.

For microbial samples, nodule data was obtained by collecting and counting all nodules in the root system of each library plant. Nodules were washed with 6% NaClO, pulverized with sterile pestles, vortexed for 30 s and then rinsed three times with 1 mL of sterile PBS. Samples were then centrifuged for 10 min at $400g$ to enrich the supernatant for undifferentiated cells and the pellet for bacteroids. Undifferentiated nodule bacteria primarily reside within the supernatant after centrifugation, while bacteroids live intracellularly within plant cells and predominantly concentrate in the pellet with the plant matter. DNA was extracted from each group of samples using a Qiagen Plant Pro kit, with the lysis buffer supplemented with 50 µL (50 mg/mL) lysozyme, 20 µL (20 mg/mL) proteinase K, and 5 µL (100 mg/mL) RNase A and incubated for 30 min at 37 °C to improve cell lysis and DNA quantity/quality. The starting "time0" library samples were processed using a QiaGen Powersoil kit with a supplemented lysis buffer.

Microbial DNA was amplified using a "BarSeq" primer set to recognize the barcode insertion site and the surrounding inactivated gene. These primers were truseq-aligned and automatically demultiplexable by Illumina software, allowing all 45 samples to be sequenced on a single lane. PCR conditions were set as described previously [1] and verified by gel electrophoresis before further processing. PCR products were cleaned up using AMPure XP Bead solution with a 1.2x ratio to select for DNA fragments  100 bp in size, pooled together with the Zymo Clean and Concentrator kit, and then pooled BarSeq product was sent to IU Center for Genomics and Bioinformatics for Illumina sequencing on a Nextseq 2k.

#### 1.4 Plant and microbial data

With plant phenotypic data (i.e., plant biomass, height, chlorophyll content, and leaf count), we tested for the effect of nitrogen enrichment using Welch's $t$-tests and ANOVA (Fig.S1). Plant biomass was based on shoot biomass pooled per-pot and measured after drying.

To generate gene fitness scores from sequencing data, we used the software suite for FEBA (Fitness Encyclopedia of Bacteria and Archaea) [1, 2]. FEBA takes as input the raw reads of an Illumina sequencing run, demultiplexed using Illumina software (for this primer set). These data were processed by the IU CGB and received in raw form as compressed fastq.gz files. The mutant library was created previously for a series of experiments by the Deutschbauer lab [2]. The Deutschbauer lab provided an aliquot of Sm1021 RB-TnSeq library for use in this experiment, and preliminary analyses which defined the mutant pool were obtained from the FEBA database. These analyses mapped barcode insertion locations to genome loci and screened ”useless” barcodes (those inserted in the first or last 10% of a gene or into an intergenic region) out of the pool.

Our data analysis started from the "BarSeq" step as outlined elsewhere [2]. The script MultiCodes.pl scanned the Illumina sequencing reads from our experiment and counted all the recovered barcodes. CombineBarseq.pl then merged the barcode counts with the "pool" file created by DesignRandomPool.pl. BarSeqR.pl took the resulting table, merged BarSeq outputs for each gene for each treatment group, and calculated gene-level fitness per gene per group by taking the $log_2$-ratios between endpoint barcode relative abundances and ”time0” relative abundances from the starting library grown in rich media [1, 2]. Subsequent analyses were carried out using tools beyond the FEBA pipeline. Only samples with more than 1 million sequencing reads were included in these subsequent analyses.

#### 1.5 Statistical analysis of gene fitness data

To assess overall survivorship and recovery of barcodes, the gene fitness data from each treatment group was pooled and analyzed for Shannon diversity, evenness, and richness (Table.S1). These metrics describe the overall size and distribution of recovered barcode counts across mutants in each environment. These results demonstrate reasonable coverage for each replicate. This was also used to verify that the starting (”time0”) library pool performed as expected and would serve as a viable point of comparison for the selective pressures experienced in the experimental treatments. Indeed, all or nearly all recoverable barcodes were present in the time0 sequencing data, and evenness was uniformly very high (indicating little to no selection).

#### 1.6 Functional genomics analyses

Functional annotation of the *S meliloti* genome, as required for downstream analyses, was conducted by matching old gene accession ids (format: SmX00000) to new ids (format: SM_RS00000) to attain maximum coverage. We then passed the updated genome to eggNOG-mapper to obtain COG and KEGG annotations as well as amino-acid sequences. The annotated genome was then partitioned by essentiality for symbiosis by conducting a Reciprocal Best-Hits (RBH) analysis between amino-acid sequences of the *S. meliloti* genome and the 603 genetic regions as determined elsewhere [3] to be essential for nodulation in *Rhizobium leguminosarum* Rlv3841 as it colonizes pea roots. This yielded a set of 832 genes taken to be essential for symbiosis in *S meliloti*.

To analyze variation in gene fitness across sites, we performed unscaled PERMANOVAs [4] to analyze how gene fitness varied by nitrogen treatment, as well as two-sample Kolmogorov-Smirnov tests of ranked gene fitness for each possible treatment or cell-type comparison. To visualize the distribution of ranks used for these tests, "rank-fitness" curves were generated by plotting fitness rankings (averaged over replicates) consecutively against the gene-level fitness scores for each mutant in the pool. These plots show how the distribution of fitness effects differs between low nitrogen and high nitrogen bacteroids as well as between the experimental and time0 groups (Fig. S2).

Fitness data were further parsed using gene set enrichment analysis to determine particular metabolic pathways and functions driving the fitness variation seen across nitrogen treatments. This was accomplished using $fGSEA$, an R package designed to run gene set enrichment analyses efficiently on large datasets [5]. Genes were first ranked according to the modified $t$-statistic tabulated by the FEBA software, which accounts for effect size and significance of each gene, by averaging scores for each gene within each nitrogen treatment and cell-type group. fGSEAs were then run for both COG categories and KEGG pathways to capture selection for general metabolic functions (e.g., carbohydrate/nucleotide metabolism) as well as pathway-level details, respectively.

Analyses were separately conducted for genes essential to symbiosis and genes nonessential to symbiosis as determined by the RBH analysis. However, all significant results depicted in Figs. 1 and 2 occurred among essential symbiosis genes — except for "Val, Ile, Leu degradation" genes, which were nonessential, being detrimental to fitness at high nitrogen among undifferentiated nodule bacteria. This particular gene function has been shown to be important for nodule symbiosis in some rhizobia-legume systems; it is possible that our RBH method mis-annotated this function as nonessential for symbiosis. As such, it is displayed along with results for symbiosis genes.

Gene set enrichment analysis conducts a random walk down a list of genes ranked by modified $t$-statistic, adding points to a running score when encountering a gene that belongs to a query set and subtracting points when the gene does not belong to the set of interest. The maximum (if positive) or minimum (if negative) value obtained during this process is the enrichment score, which is converted to a normalized enrichment score (NES) to enable comparisons across multiple GSEAs [5]. Genes that are significantly concentrated toward the top or bottom of the ranking accumulate high and low enrichment scores and are flagged as "enriched" or "depleted", respectively. The genes that contribute to this maximum or minimum score are most responsible for the overall enrichment of that set and are referred to as leading-edge genes [5]. Significant fGSEA results were plotted to visualize functions and metabolic pathways most beneficial/detrimental to knockout mutant fitness at each nitrogen level, for each cell-type, for both symbiosis and non-symbiosis genes. This allowed identification of specific functions under selection at each nitrogen level, while also isolating shared phenotypes that may correspond to the general demands of the plant-associated rhizosphere environment and/or metabolic pressures specific to within-nodule growth. Leading-edge genes for each treatment group were collated and reported (Table. S2). Furthermore, to assess the distribution of genes under selection across replicons, a Chi-square analysis was performed comparing the number of leading-edge genes on chromosome vs. plasmids to the total number of genes on chromosome vs. plasmids.

While high-attrition, bottlenecked environments present challenges for the use of RB-TnSeq, we demonstrate that existing methods for parsing other kinds of genome-wide $log_2$-ratio data can be applied to RB-TnSeq data with some caveats. Fold-changes below |2| may be more biologically meaningful when quantifying barcoded mutant survivorship rather than transcription, as fluctuations in transcript abundance are often incidental whereas mutant survivorship directly measures mutant fitness. Gene fitness is inversely proportional to knockout mutant fitness, as a mutant which outperforms WT has been unburdened of a detrimental gene while one which underperforms WT has been deprived of something useful. To prevent confusion, the results of these analyses are sign-inverted so as to depict gene fitness rather than mutant fitness.

Improving the performance of this technique to capture subtle differences between complex environments could enable high-throughput genotype-phenotype mapping across complex environments with highly multidimensional fitness landscapes. Currently, differences between complex environments can be difficult to capture when both conditions differ strongly and similarly from the rich media in which the time0 group is maintained. Future RB-TnSeq experiments could make finer-grained inquiries into these environments by comparing experimental groups to a time0 baseline grown in defined media tailored to the environment of interest [6].

##### 1.7 Data availability

All code used in this study is available at https://github.com/LennonLab/rhizo.rb.tnseq. RB-TnSeq data analysis scripts can be found on the FEBA codebase, and mutant pool information and results from previous RB-TnSeq experiments [1, 2] can be found here.

#### 2 Supplementary Figures and Tables

| ID | Pot | N enrichment | Cell-type | Diversity | Evenness | Richness | %rec |
|---|---|---|---|---|---|---|---|
| X21 | 1 | yes | Nodule | 2.32 | 5.76E-06 | 3640 | 70% |
| X22 | 2 | yes | Nodule | 2.01 | 8.75E-06 | 3115 | 60% |
| X23 | 3 | yes | Nodule | 3.06 | 1.43E-04 | 646 | 12% |
| X24 | 4 | yes | Nodule | 1.47 | 1.54E-05 | 2508 | 48% |
| X26 | 6 | no | Nodule | 1.48 | 6.35E-06 | 3762 | 73% |
| X27 | 7 | no | Nodule | 0.33 | 7.09E-05 | 3505 | 68% |
| X28 | 8 | no | Nodule | 2.48 | 6.30E-06 | 3211 | 62% |
| X29 | 9 | no | Nodule | 3.55 | 6.95E-05 | 2368 | 46% |

| ID | Pot | N enrichment | Cell-type | Diversity | Evenness | Richness | %rec |
|---|---|---|---|---|---|---|---|
| X30 | 10 | no | Nodule | 1.94 | 1.12E-05 | 2548 | 49% |
| X31 | 1 | yes | Bacteroid | 1.38 | 6.30E-05 | 971 | 19% |
| X32 | 2 | yes | Bacteroid | 1.60 | 1.03E-05 | 3083 | 60% |
| X33 | 3 | yes | Bacteroid | 3.06 | 3.94E-05 | 2963 | 57% |
| X35 | 5 | yes | Bacteroid | 2.84 | 1.36E-04 | 1541 | 30% |
| X37 | 7 | no | Bacteroid | 0.73 | 6.01E-05 | 3516 | 68% |
| X40 | 10 | no | Bacteroid | 1.95 | 7.60E-06 | 3757 | 73% |
| X41 | NA | NA | time0 | 7.94 | 1.71E-02 | 5175 | 100% |
| X42 | NA | NA | time0 | 7.91 | 1.78E-02 | 5175 | 100% |
| X43 | NA | NA | time0 | 7.86 | 1.63E-02 | 5166 | 99% |
| X44 | NA | NA | time0 | 7.92 | 1.69E-02 | 5175 | 100% |
| X45 | NA | NA | time0 | 7.95 | 1.74E-02 | 5175 | 100% |

**Table 1. Barcode recovery statistics for Sm1021 mutant pool.** Metadata describing the pool of mutants recovered at the end of the 8-week experimental period. Each replicate is described by nitrogen level, cell-type, and pot number. The recovered pool of mutants was characterized at a broad level by measuring the Shannon diversity, normalized median evenness, and richness. Richness data are also presented as %recovered, which is the richness number divided by 5175, the total number of recoverable genes in the library genome.

Download Supplementary Table S2

**Table 2. Genes under selection by treatment group.** This table lists genes under selection in each treatment group as determined by our gene-set enrichment analyses, identifying nitrogen-dependent genes as well as genes which experienced selection at both low and high N for each cell-type. The GSEA algorithm determines which gene sets are enriched or depleted, but it also provides a list of ”leading-edge genes” for each significant set. These are simply the genes which appear significantly close to the top or bottom of the genome-wide fitness ranking and contribute to the enrichment score. GSEAs are run on a treatment-specific basis and therefore can identify genes and functions specific to each individual replicate group. These are presented here along with locus tags, gene names, COG categories, essentiality for symbiosis (as determined by our RBH analysis), KEGG annotations, and gene descriptions. Only eight genes were significantly enriched or depleted among the low-nitrogen bacteroids; all of these were also significant at high nitrogen, along with many other genes exclusively under selection at high nitrogen. Among the undifferentiated nodule bacteria, a handful of genes experienced selection only at low nitrogen, a large number experienced selection at both low and high nitrogen (the ”nodule common genes” in the table), and another set experienced selection only at high nitrogen.

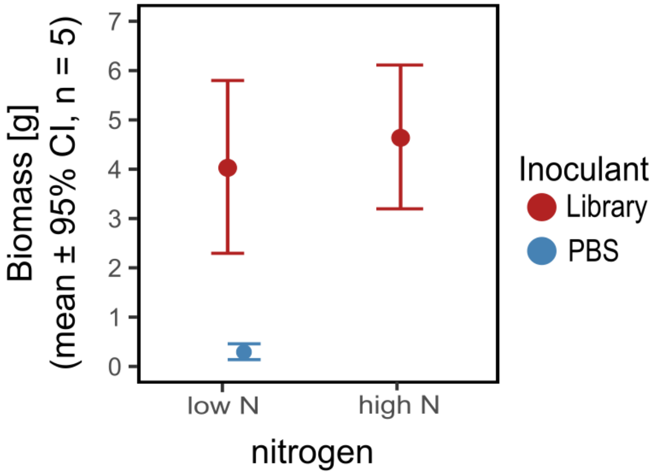


**Fig 1. Mean plant biomass by treatment group.** Plant biomass was not significantly affected by soil nitrogen enrichment. Interval plot shows mean dry biomass in grams, ± 95%. CI, of *M. sativa* host plants after 8 weeks of growth post-inoculation with *Ensifer meliloti* RB-TnSeq library (red) or PBS control (blue). Measurements were taken on above-ground biomass after plants oven-dried for 72 h at 80 °C.

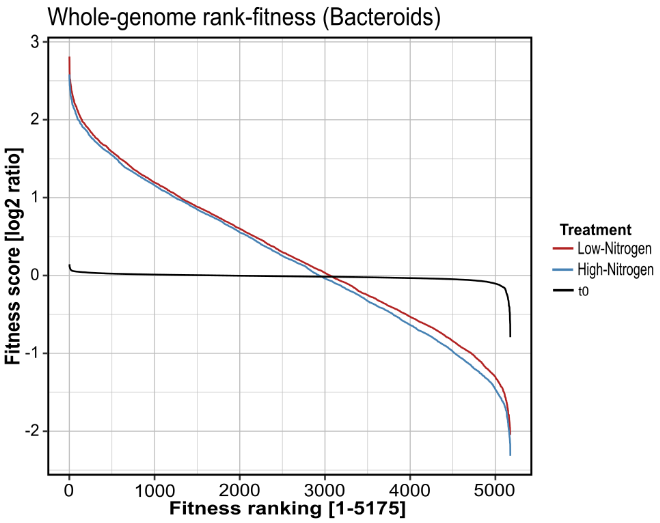


**Fig 2. Distribution of fitness effects at low vs high nitrogen for bacteroids.** Rank-fitness plots depicting gene fitness ranking against gene fitness score as calculated by the FEBA software. Fitness scores are $log_2$ ratios of barcode relative abundances after growth compared with the starting library, and rank-order is obtained by arranging these from greatest to least. The distributions of fitness effects across every gene in the genome for high nitrogen bacteroids (blue), low nitrogen bacteroids (red), and the time0 library (black). K-S tests revealed significant difference in the distribution of fitness effects between low-nitrogen and high-nitrogen bacteroids, while the time0 library is nearly flat as expected for a library undergoing no significant selection.

3. Poole P, Ramachandran V, Terpolilli J. Rhizobia: from saprophytes to endosymbionts. Nat Rev Microbiol. 2018;16(5):291–303.

4. Oksanen J, Simpson GL, Blanchet FG, Kindt R, Legendre P, Minchin PR, et al..

vegan: Community Ecology Package; 2022. Available from: https://CRAN.R-project.org/package=vegan.

5. Korotkevich G, Sukhov V, Budin N, Shpak B, Artyomov MN, Sergushichev A. Fast gene set enrichment analysis; 2016.

6. Borchert AJ, Bleem A, Beckham GT. Experimental and analytical approaches for improving the resolution of randomly barcoded transposon insertion sequencing (RB-TnSeq) studies. ACS Synth Biol. 2022;11(6):2015–2021.
